# A Method for Calculating the Least Mutated Sequence in DNA Alignment Based on Point Mutation Sites

**DOI:** 10.1101/2023.11.14.567125

**Authors:** Libing Shen

**Affiliations:** International Human Phenome Institutes (Shanghai), Shanghai, 200433, P.R. China; Greaeter Bay Area Institute of Precision Medicine, Guangzhou, 511462, P.R. China

## Abstract

It is hard to decide the root of a phylogenetic tree when no suitable outgroup sequence is available, especially in a high mutation rate system. Here we propose a bioinformatics method for calculating the least mutated sequence in DNA alignment based on point mutation sites. By parsimony principle, the least mutated sequence should be the ancestor (phylogenetic root) of the other sequences in an alignment result. The intuition suggests that our method would be sound only if enough sequences were collected. In this study, we used DNA mutation simulation data to test our method. The simulation results show that our method is very reliable and just a small proportion of sequences is required to find the least mutated sequence when the sampling is random. When the ancestral sequence is present in the dataset, the least mutated sequence calculated by our method is the ancestral sequence itself, i.e., the phylogenetic root of the dataset. When the ancestral sequence is not in the dataset, the least mutated sequence calculated by our method is the one closest to the phylogenetic root. Furthermore, the simulation demonstrates that our method is robust against reverse mutation and saturation mutation. The performance of our method improves with the increasing number of sequences. We developed the program based on this method which also calculates the transition/transversion ratio for each sequence in an DNA alignment. It can be used as a rough measure for estimating selection pressure and evolutionary stability for a sequence. In conclusion, our method and the corresponding programs could be a useful auxiliary tool for bioinformaticians and evolutionary biologists.

## Introduction

Phylogenetic analysis remains to be the most fundamental part in the research fields of bioinformatics and evolutionary biology. Now due to the advancement of sequencing technology, molecular sequence data become the absolute base for biological phylogeny. DNA and RNA have four different nucleotides while protein commonly has 20 different amino acids. This fact renders protein more robust than DNA or RNA in phylogenetic analysis. Two protein sequences can be correctly aligned even if they share only 25% sequence identity (1, 2), whereas the correct alignment of two DNA sequences requires that they must share more than 50% sequence identity (3). Thus, protein sequences are more advantages in performing the phylogenetic analysis for the biological entities with long generation time, i.e., metazoan; nucleotide sequences are more suitable for the phylogenetic analysis of the biological entities with short generation time, i.e., bacteria or virus.

Short generation time is always accompanied with high mutation rate. In a high mutation rate system, one common problem is how to root its phylogenetic tree, especially when there is no proper outgroup. Using a distantly related sequence as the outgroup will create a problem called long-branch attraction in phylogenetic analysis (4). The fast-evolving sequences (long branch) on the phylogenetic tree would be wrongfully placed close to the outgroup sequence as if they were closely related. It would lead to the wrong evolutionary relationship between fast-evolving operational taxonomic unit and outgroup operational taxonomic unit. This phenomenon is worse in the phylogenetic tree based on DNA alignment than on protein alignment. Nevertheless, DNA sequences are more useful than protein sequences in evolutionary analysis such as the calculation of synonymous and nonsynonymous substitutions, which are commonly used to estimate the evolutionary pressure among operational taxonomic units. This practice also requires a correct phylogenetic root. It serves as the sequence template for documenting the observed mutations.

In this paper, we propose a bioinformatics method for searching the least mutated sequence in DNA alignment based on point mutation sites. By parsimony principle, the least mutated sequence should be the phylogenetic root for all the other sequences in an alignment result. Our method is a non-parameter method and uses the point mutation sites in a DNA alignment result for calculation. It doesn’t need any outgroup sequence as reference. We applied our method to the simulation dataset based on real viral sequence for testing. The testing results show: 1) our method only needs a very small proportion of sequences to find the root sequence under random sampling; 2) our method is quite robust against reverse mutation and saturation mutation, whose accuracy rises with the increasing number of sampling sequences. Finally, we found that reconstructing a phylogenetic tree could be used to verify the result of our method.

## Data and Method

### Global pair-wise point mutation calculation

In a correct DNA alignment, each nucleotide site is a positional homology, which means the nucleotides at this site are descended from a common ancestral nucleotide (a common ancestor). If a point mutation happened to one sequence in the alignment, the mutated nucleotide would be different from the nucleotides of the other sequences at this site. The site becomes a mutation site which harbors more than one type of nucleotide. If each sequence is pair-wise compared to the other sequences in the alignment, the number of global point mutations can be calculated for each sequence. By parsimony principle, the sequence that has the least number of point mutations should be the phylogenetic root in the alignment (Figure 1).

**Figure 1.**
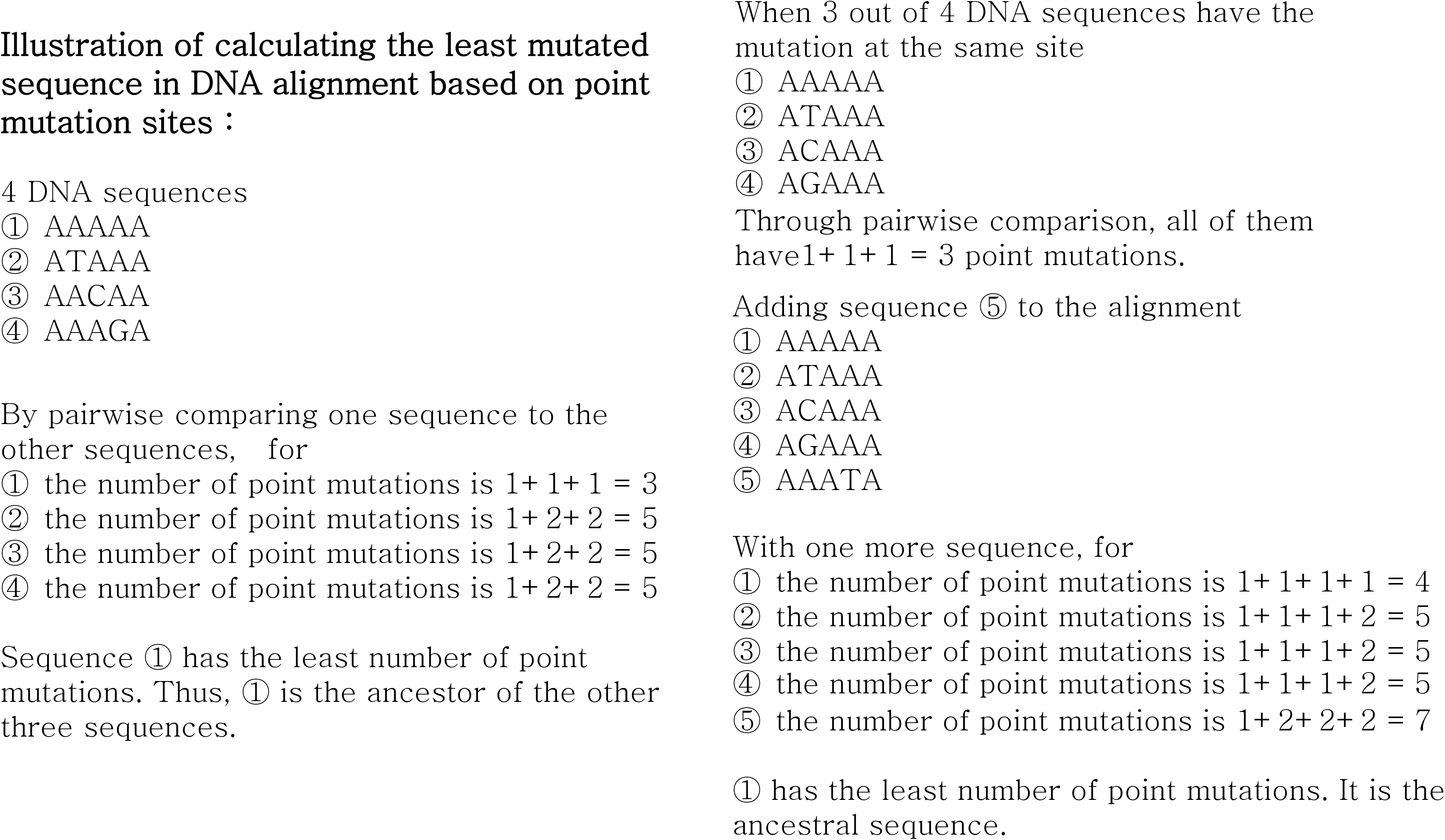
The illustration how to use the point mutation sites in a DNA alignment to calculate the number of mutations for each sequence and deduce the least mutated sequence.

Sometimes a nucleotide site can be a mutation hot spot where multiple point mutations or reverse mutation occurred. If the number of sequences is limited in the alignment, it will create the difficulty for our method. We noticed that such difficulty could be solved by adding additional sequence to the alignment (Figure 1). Thus, this method intuitively seems to work better with more sequences in the alignment.

### Implementation

We implemented this method by two algorithmic steps. First, each point mutation site is extracted from the alignment file and then all point mutation sites are organized into a matrix file. Second, each nucleotide in the matrix is compared with the other nucleotide at the same site in a pair-wise comparison fashion and the number of mutations is documented. The number of mutations at each site is summed for each sequence in order to calculate global point mutations. Finally, each sequence will be sorted according to its number of point mutations from the smallest to the largest. The first sequence is the least mutated sequence. The number of transition and transversion mutations are also calculated in the second step. Thus, the transition/transversion ratio is provided for each sequence in the final result.

### Simulation data for testing our method

We used the complete coding genome of SARS coronavirus (AY274119.3) as the ancestral sequence to generate the simulation data for our test. It has 29124 nucleotides in length. The simulation is as follows: 1) we gradually generate 1 to 200 random point mutations in the ancestral sequence in order to simulate the natural mutation process; 2) the simulated natural mutation process will be repeated 100 times to produce 20,000 descendant mutated sequences; 3) we randomly select 9, 99 or 499 descendant mutated sequences and put back the ancestral sequence to constitute a complete test dataset with total 10, 100 or 500 sequences; 4) 1000 simulation datasets comprising 10, 100 or 500 sequences (total 3000 simulation datasets) are generated for testing whether the ancestral sequence is the least mutated one. We also generate 3000 simulation datasets without putting back the ancestral sequence. In the third step, we just randomly select 10, 100 or 500 descendant mutated sequences to constitute a complete test dataset.

Finally, we tested the performance of our method under the conditions of reverse mutation and saturation mutation. We extracted a 100-nucleotide fragment from the coding region of SARS coronavirus (AY274119.3) to repeat the simulation process above. Because we gradually generate 1 to 200 random point mutations in a 100-nucleotide long sequence, there must exist the circumstances of reverse mutation and multiple mutations at the same nucleotide site in this simulation.

## Result

### The performance of our method in the simulation of natural mutation process

Table 1 shows the performance of our method in the simulation of natural mutation process. When the ancestral sequence is in the test dataset, our method can 100% correctly find the least mutated sequences in all three different sample size datasets. When the ancestral sequence is not put back in the test dataset, our method exhibits a slightly different accuracy in different sample size dataset. Our method had 97.6% accuracy in the test dataset with 10 randomly selected sequences whereas it achieved 99.7% and 99.9% accuracy in the test dataset with 100 or 500 randomly selected sequences.

**Table 1.**
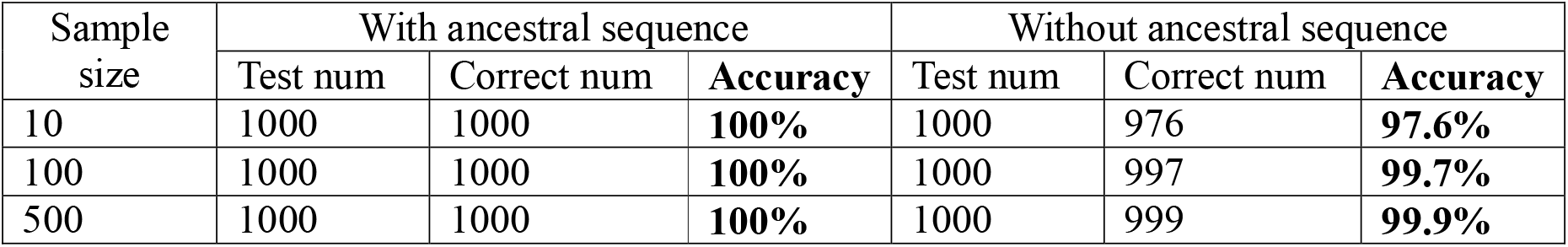
Test of our method using the simulation of natural mutation process. The length of the ancestral sequence is 29124 nucleotides.

### The performance of our method in the simulation of reverse mutation and saturation mutation

Table 2 shows the performance of our method in the simulation of reverse mutation and saturation mutation. When the ancestral sequence is in the test dataset, our method shows an above 95% overall accuracy in all three different sample size datasets. It achieved 99% accuracy in the test dataset with 100 or 500 randomly selected sequences. When the ancestral sequence is not put back in the test dataset, our method exhibits a varying accuracy across different sample size datasets. Its accuracy increases with the sample size. It had an 82.7% accuracy in the dataset with 10 sequences while had an 93.6% accuracy in the dataset with 100 sequences. Its accuracy rises to 97.5% when the test datasets have 500 sequences.

**Table 2.**
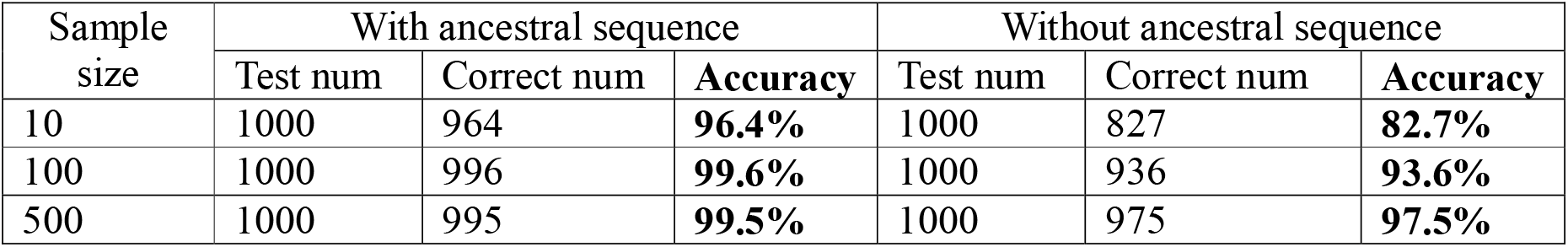
Test of our method using the simulation of reverse mutation and saturation mutagenesis. The length of the ancestral sequence is 100 nucleotides.

## Discussion

Our method shows a reliable and robust performance in the simulation tests of natural mutation process, especially without the presence of reverse mutation and saturation mutation. In all 3000 simulation tests which has the ancestral sequence in the dataset, our method 100% correctly calculated the ancestral sequence as the least mutated sequence. In the simulation tests which don’t include the ancestral sequence in the dataset, our method demonstrates a stable performance as well. It achieved 97.6% accuracy in the simulations with the sample size of 10 sequences and achieved above 99% accuracy in the simulations with large sample sizes.

In a sequence alignment result, the least mutated sequence is not necessarily the ancestral sequence (the phylogenetic root). If the ancestral sequence is not present in the dataset, the least mutated sequence is the one evolutionarily closest to it in an alignment result. We found that one could construct a phylogenetic tree to distinguish whether the least mutated sequence is the phylogenetic root or not. When an unrooted tree is displayed in a radiation fashion, the ancestral sequence (the phylogenetic root) is in the right center of the tree (Figure 2). If the mutated sequence is not the ancestral sequence, it will be the closest taxon to the right center of the tree (Figure 3). We further found that phylogenetic tree could be used to examine whether the result of our method is correct or not. Figure 4 shows an erroneous result from our simulation test. In this case, the least mutated sequence calculated by our method is AY274119.3_iter:94_mut:67. However, the true least mutated sequence should be AY274119.3_iter:22_mut:62 in this alignment. Compared with the ancestral sequence, the former has 67 point mutations and the latter has 62 point mutations. The reason why our method produced a wrong least mutated sequence is that this test dataset has a sequence of AY274119.3_iter:94_mut:148. It is a derivative sequence from AY274119.3_iter:94_mut:67. Both of them are on the same evolutionary branch. Its existence renders AY274119.3_iter:94_mut:67 have 67 point mutations deducted from its final calculation, which leads our method to a wrong result. This case also shows that adding more sequences to the final dataset could avoid this kind of error and increase the accuracy of our method.

**Figure 2.**
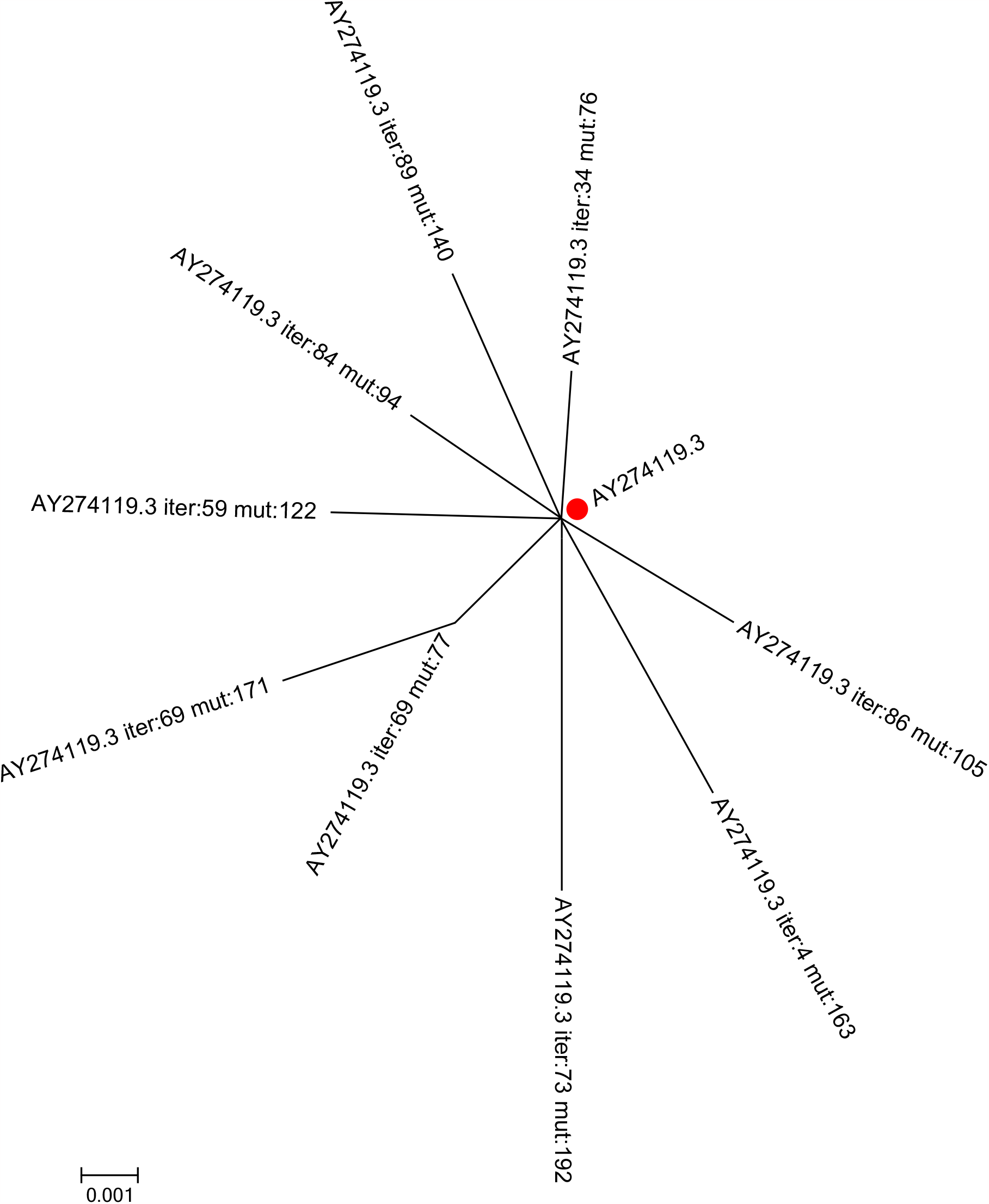
Neighbor-joining tree for the dataset including the ancestral sequence. When the least mutated sequence is the ancestral sequence, its position is in the center of the phylogenetic tree. Red filled circle 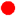 indicates the least mutated sequence.

**Figure 3.**
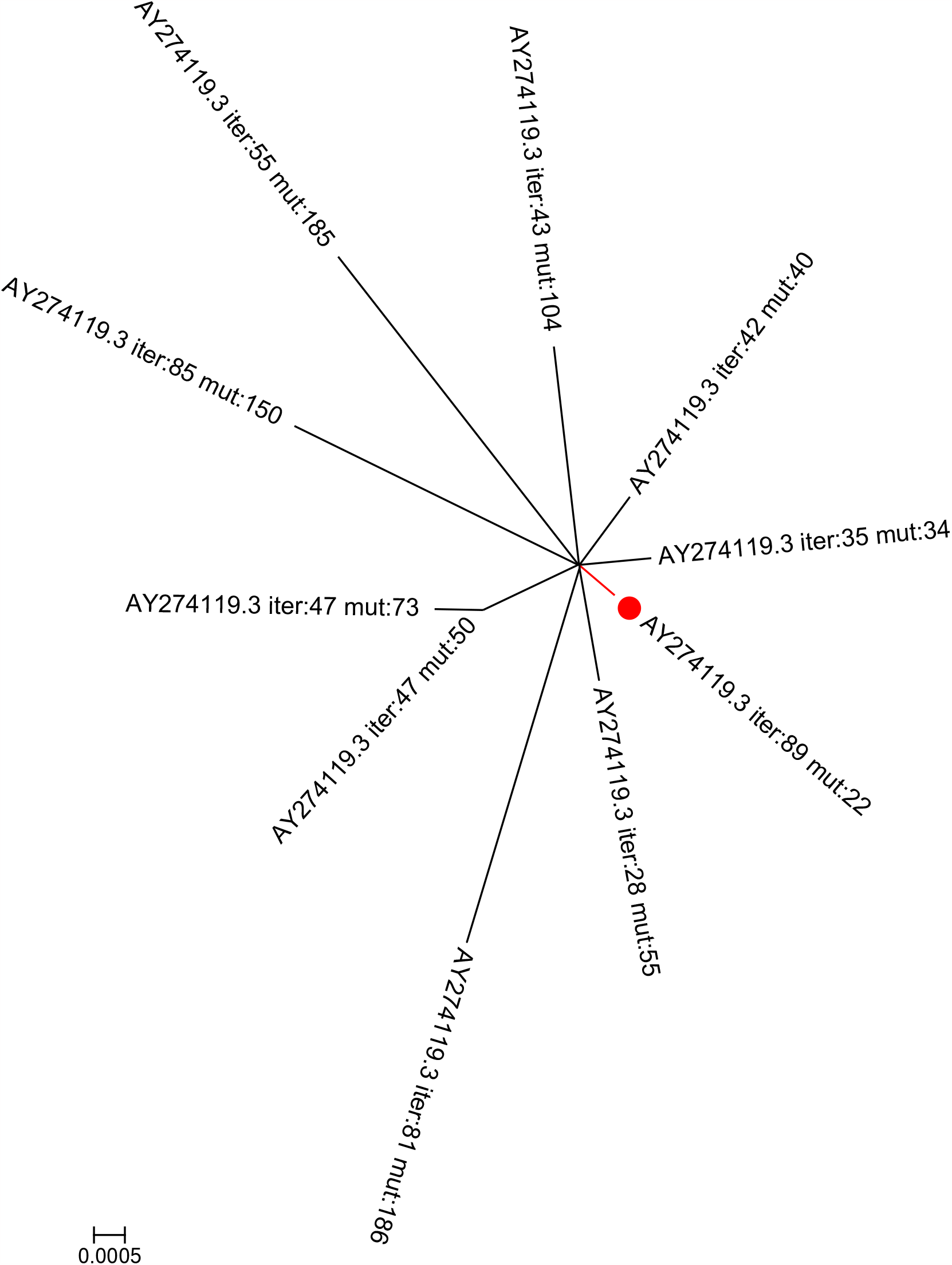
Neighbor-joining tree for the dataset not including the ancestral sequence. When the least mutated sequence is not the ancestral sequence, its position is closest to the center of the phylogenetic tree. Red filled circle 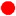 indicates the least mutated sequence.

**Figure 4.**
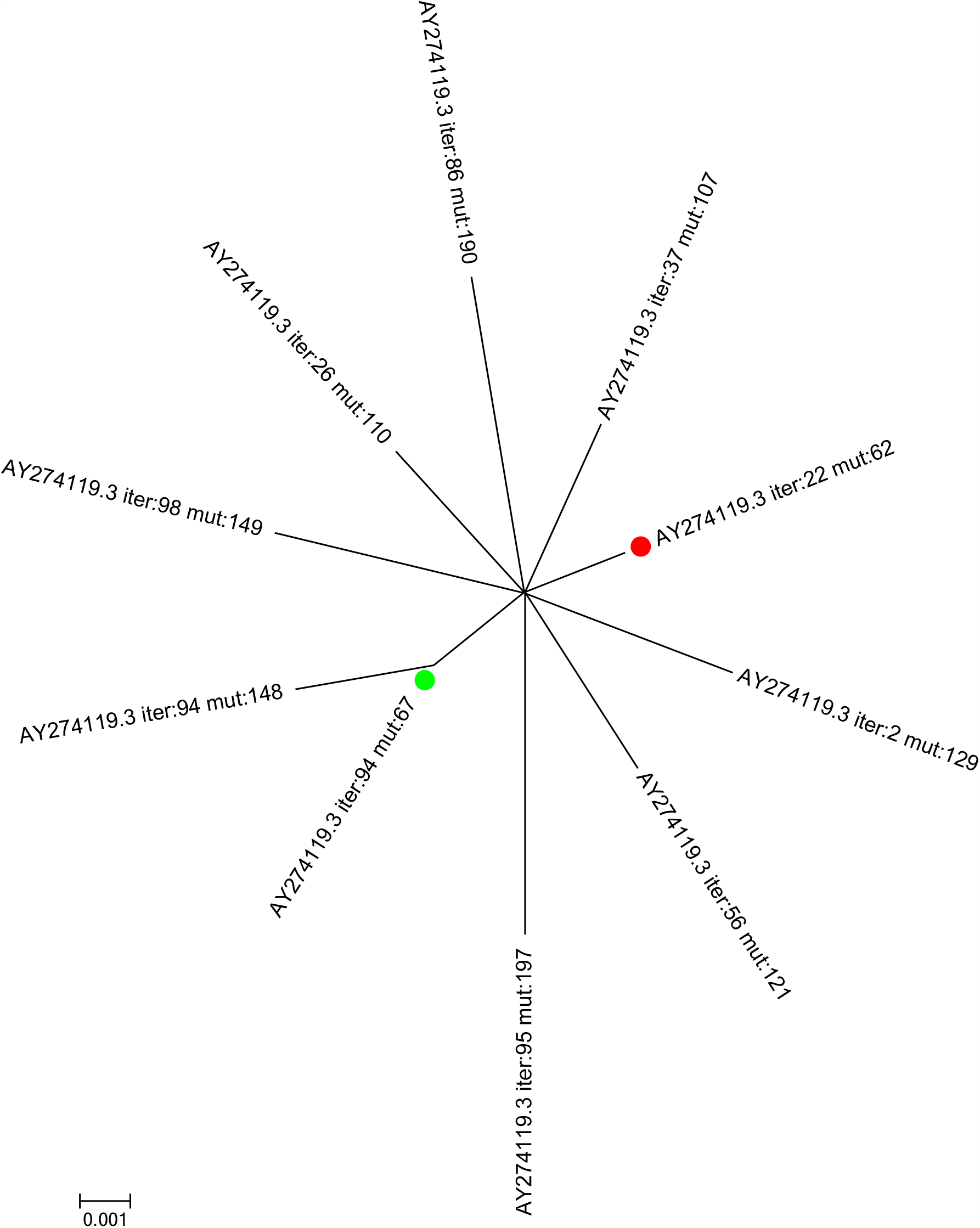
Neighbor-joining tree shows an example of our method’s error. Green filled circle 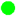 indicates the least mutated sequence calculated by our method. Red filled circle 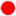 indicates the actual least mutated sequence in this dataset.

Reverse mutation and saturation mutation are big troubles for any phylogenetic inference methods (5, 6). They would blur the true evolutionary history of a DNA sequence. Our simulation result shows that our method could reach a quite reasonable accuracy even with the great presence of reverse mutation and saturation mutation. In the real world, it is almost impossible to a perfect dataset including all mutated sequences. The number of sampled sequences is always far less the number of total sample size. In our simulation tests, we generated 20,000 mutated sequences and the largest sample size is 500 sequences which is only 2.5% of total sample size (500/2000). Increasing the number of sequences in evolutionary analysis can often make the real evolutionary process clearer and improve the accuracy of our method. For the same reason, when the ancestral sequence is included in the dataset, our method can deliver a better performance. Thus, we propose that when using our method collect DNA sequences as many as possible. The more sequences you collect, the more likely you will include the ancestral sequence in your data.

## Resource availability

The corresponding programs for this study can be downloaded from https://github.com/libing-shen/Least-mutated-sequence. The simulation program and results for this study can be downloaded from https://github.com/libing-shen/Least-mutated-sequence/tree/main/simulation.

## Compliance and ethics

All authors declare no potential competing interest. This work used the virtual data only and ethical approval is not applicable.

## Authors’ contributions

LBS designed and wrote the Perl programs, analyzed the simulation results, and wrote the manuscript.

## Consent for publication

The author consented the right for publication.

